# The short-term high fat diet-induced increase in %5-methylcytosine expression in peripheral blood T lymphocytes, is attenuated by low-dose aspirin

**DOI:** 10.1101/825125

**Authors:** Tinashe Mutize, Phiwayinkosi V. Dludla, Zibusiso Mkandla, Bongani B. Nkambule

## Abstract

**Objective:** To assess peripheral lymphocyte DNA methylation profiles in prediabetes using a high fat-diet-fed C57BL/6 animal model. We further evaluated whether low dose-aspirin, or low-dose aspirin in combination with metformin, could modulate global DNA methylation levels in peripheral blood lymphocytes.

**Methods:** Twenty-eight (28) male C57BL/6 mice were used in two experimental phases. The first experiment involved animals (n=16) which were randomised to receive a low-fat diet (LFD) or high-fat diet (HFD) (n = 8/group) for 10 weeks. Whereas in the second experiment, HFD-fed mice (n=15) were randomised into 3 treatment groups, a low-dose aspirin (LDA), LDA and metformin group, and a clopidogrel group. DNA methylation profiles of were determined using flow cytometry.

**Results:** The HFD group showed moderate weight gain and elevated postprandial blood glucose levels when compared to the LFD group after 2 weeks of HFD-feeding (p < 0.05). Interestingly, the HFD group had elevated levels of T cells expressing high levels %5-methylcytosine (p<0, 05). Notably, these elevated levels were lowered by short-term low-dose aspirin treatment.

**Discussion:** T cells are involved in the propagation of the inflammatory response. Persistent T cell activation promotes chronic inflammation and insulin resistance. Low-dose aspirin may be effective in modulating T cell-specific global methylation.

## Introduction

The global prevalence of type 2 diabetes (T2DM) continues to increase with several studies linking aberrant DNA methylation with the development of these metabolic disease (Van Otterdijk et al., 2017; Zhang et al., 2017; Shah et al., 2019; Wittenbecher et al., 2019). Aberrant DNA methylation profiles are associated with an increased risk of developing T2DM (Zhou et al., 2018; Zhang et al., 2018). Differential DNA methylation has been reported in patients with T2DM (Kuroda et al., 2009; Nilsson et al., 2014; Dayeh et al., 2014; Zhang et al., 2018). DNA hypermethylation (Ban et al., 2002; Yang et al., 2011; Gu et al., 2013; Zou et al., 2013; Seman et al., 2015; Bacos et al., 2016) and DNA hypomethylation (Zhang et al., 2014; Nilsson et al., 2015) levels have been reported in T2DM. Moreover, increased levels of global DNA methylation were associated with insulin resistance (Zhao et al., 2012) and in another study, aberrant DNA methylation was reported in women with obesity-related systemic insulin resistance (Zhang et al., 2018). Hypermethylation on the insulin promoter in type 2 diabetic patients is also associated with the downregulation of the insulin gene in human pancreatic islets (Yang et al., 2011). Furthermore, Kuroda *et al.*, also reported that the expression of the insulin gene was regulated by differential DNA methylation (Kuroda et al., 2009).

Several studies have associated dietary patterns with DNA methylation profiles studies and obesity-related conditions such as chronic inflammation, insulin resistance, T2DM and subsequently cardiovascular disease (Mckay & Mathers, 2011; Park et al., 2017; Milagro et al., 2013).

Chronic inflammation promotes insulin resistance and the development of type 2 diabetes (Goldfine et al., 2011). Chronic inflammation also provides a link between type 2 diabetes and cardiovascular disease (Goldfine et al., 2011). Previous studies have shown that differential methylation in adipose tissues, muscles and pancreatic islets are associated with localised inflammation in patients with T2DM (Kulkarni et al., 2012; Yang et al., 2012; Ribel-Madsen et al., 2012; Nilsson et al., 2014; Dayeh et al., 2014).

Low-dose aspirin (LDA) is widely used for the primary prevention of cardiovascular events in patients with type 2 diabetes who are at an increased risk of developing thrombotic complications (Goldfine et al., 2011). Furthermore, the use of low-dose aspirin as a primary preventive measure for patients with type 2 diabetes and heart failure (HF) has been reported to be associated with lower all-cause mortality, though in the absence of other contraindications including a history of myocardial infarction, stroke, coronary artery disease, peripheral artery disease (Abi Khalil et al., 2018) (Chang et al., 2013). Furthermore, it has been suggested that an increased dose of aspirin or twice-daily low-dose aspirin therapy could be possible therapeutic options for cardiovascular prevention in diabetes mellitus patients (Capodanno & Angiolillo, 2016). The combination of aspirin with a potent anti-platelet drug such as clopidogrel is an antiplatelet therapy option of choice in the prevention of macrovascular conditions (Silber et al., 2015; Smith et al., 2006; Anderson et al., 2007; King et al., 2008; Werf et al., 2008; Kushner et al., 2009). Notably, the occurrence of cardiovascular events in coronary artery disease patients receiving dual clopidogrel and low-dose aspirin therapy is associated with a poor response to clopidogrel (Kuliczkowski et al., 2009). The poor response and variability in clopidogrel action, as well as the recurrence of ischemic events in patients with stroke, has been attributed to differential DNA methylation (Gallego-Fabrega et al., 2016). In fact, hypomethylation of the ATP Binding Cassette Subfamily B Member 1 (ABCB1) gene promoter has been reported to be associated with a decreased clopidogrel response in ischemic stroke patients via increased ABCB1 mRNA expression (Yang et al., 2015).

In this study, we hypothesized that low dose aspirin (LDA) as a monotherapy and in combination with metformin (LDA + Metformin) may induce differential global DNA methylation in B and T lymphocytes (Nishimura et al., 2009; DeFuria et al., 2013). The study aimed to assess the global DNA methylation profiles of the lymphoid cell subsets using a HFD-fed animal model of prediabetes. We further assessed whether low-dose aspirin and clopidogrel modulate the global DNA methylation profiles of peripheral blood lymphocytes.

## Methods and Materials

### Animals and animal handling

Male C57BL/6 mice (n = 28) at 6 weeks of age were purchased from the Biomedical Resource Unit (BRU) at the University of KwaZulu-Natal (UKZN). The C57BL/6 mice strain is well characterized and has been shown to become glucose intolerant when kept on a high-fat diet (Pinchuk & Filipov, 2008). The mice were housed in cages at the BRU in a controlled 12-hour light/dark cycle and a temperature range of 23 - 25 °C (relative humidity: approximately 50 %). The animal well-being was monitored in accordance with the principles of laboratory animal care (National Institute of Health publication 80-23, revised 1978). The mice were allowed free access to water throughout the experimental period. The animal study followed the Animal Research: Reporting In Vivo Experiments (ARRIVE) (Kilkenny et al., 2013) (Supplementary File 1). The ARRIVE guideline was used to improve the standard of reporting in animal research. The study received ethical approval from the University of KwaZulu-Natal Animal Research Ethics Committee (AREC), under the ethics registration number AREC/086/016.

### Study design and experimental procedures

The study comprised of two major experiments (Figure 1). The first experiment comprised of 16 male mice which were randomized into two diet groups, the low-fat diet (LFD) and the high-fat diet (HFD) group (table 1). The LFD group received a low-fat diet (D12450J) (Research Diets, New Brunswick, NJ, USA) containing 10 % kcal fat, 20 % kcal Protein, 70 % kcal carbohydrates and 3.82 kcal/g energy density, whereas the HFD group (D12492) (Research Diets, New Brunswick, NJ, USA) received a high-fat diet containing 60 % kcal fat, 20 % kcal protein, 20 % kcal carbohydrates and 5.21 kcal/g energy density).

**Table 1.** Diet composition (g/kg)

| Ingredients | LFD<sup>a</sup> | HFD<sup>b</sup> |
| --- | --- | --- |
| Casein, 30 mesh | 200.00 | 200.00 |
| L-Cystine | 3.00 | 3.00 |
| Corn starch | 506.20 | - |
| Lodex 10 | 125.00 | 125.00 |
| Sucrose | 72.80 | 72.80 |
| Solka Floc, FCC200 (Fiber) | 50.00 | 50.00 |
| Soybean Oil | 25.00 | 25.00 |
| Lard | 20.00 | 245.00 |
| Mineral mix S10026B | 50.00 | 50.00 |
| Choline Bitartrate | 2.00 | 2.00 |
| Vitamin mix V10001C | 1.00 | 1.00 |
| Dye, Yellow FD&C #5, Alum. Lake 35-42% | 0.04 | - |
| Dye, Blue FD&C #1, Alum. Lake 35-42% | 0.01 | 0.05 |

<sup>a</sup> The LFD obtained from Research Diets Inc (#D12450J, rodent diet with 10% kcal% fat) provided 3.82 kcal/g from 20%, 70%, and 10% of protein, carbohydrate, and fat, respectively.
<sup>b</sup> The HFD obtained from Research Diets Inc (#D12492, rodent diet with 60% kcal% fat) provided 5.21 kcal/g from 26.2%, 26.3%, and 34.9% of protein, carbohydrate, and fat, respectively.
<sup>c</sup> Typical analysis of cholesterol in lard = 0.72 g/kg.

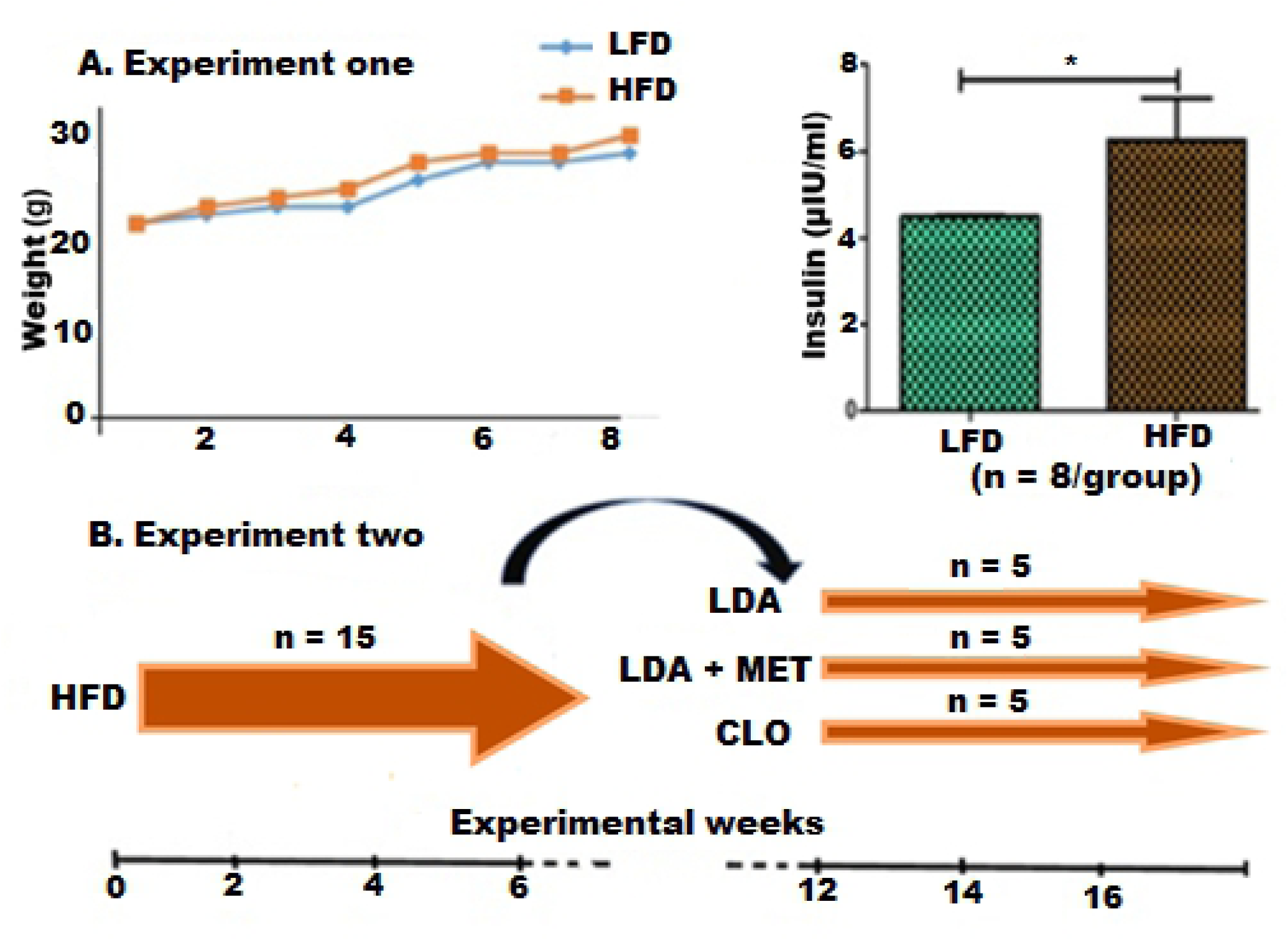

#### Experiment one

The first experiment aimed at measuring the baseline DNA methylation profiles of isolated B and T lymphocytes, following 8-weeks of high fat-diet feeding. The study comprised of 5-week old male C57BL/6 mice (n = 16). The mice were allowed a week for acclimatisation while receiving normal mice chow and had free access to water *ad libitum.* The mice were then randomised into two groups, the LFD and HFD group (n = 8/group). The animals were then housed in separate cages (n = 8/cage) based on their respective diets (figure 1). Two-hundred microliters of venous blood was drawn from each animal and baseline haematological measurements, glucose, insulin and DNA methylation measurements (%5-methylcytosine) were determined (Supplementary file 2). Monocyte and granulocytes were depleted from the whole blood samples and B and T lymphocyte were then isolated using the BD™ IMag Cell Separation system (BD Bioscience, USA) (Supplementary file 2).

#### Experiment two

In experiment two we assessed whether the short-term treatment with low-dose aspirin (LDA), LDA with metformin (LDA+MET), and clopidogrel modulates DNA methylation profiles in HFD-fed (HFF) mice. Fifteen (n=15) HFF mice were randomized into three treatment groups (n = 5/group). These included; (1) a low-dose aspirin (LDA) (3 mg/kg) (Kroesen et al., 2018) group; (2) LDA and metformin (150 mg/kg) (Saisho, 2015) group and clopidogrel (0.25 mg/kg) (Iannacone et al., 2007). The mice received their respective treatment for 6 weeks via daily oral gavage (figure 1). Blood was then drawn and the serum insulin levels, haematological indices, DNA methylation profiles were determined (Supplementary file 2).

### Statistical analysis

All statistical analysis was performed using GraphPad Prism 5 (GraphPad Software Inc.; La Jolla, CA, USA). The comparisons of baseline measurements between the two diets groups the LFD and HFD were performed using an unpaired t-test for parametric data and reported as mean and standard deviation. In addition, the one-way analysis of variance (ANOVA) was used for comparisons of DNA methylation levels across the three treatment groups, followed by the Bonferroni post-hoc test. While non-parametric data was analysed using the Mann Whitney U test and reported as median and interquartile range. For comparisons across the three treatment groups the Kruskal-Wallis test followed by the Dunn’s multiple comparison test were used. The dependent variable was the global DNA methylation level (% 5-Methylcytosine) while the independent variables were the lymphocyte subsets, treatment drug, and diet group. A p-value of < 0.05 was considered statistically significant.

## Results

### Baseline characteristics and haematological parameters following 8-weeks of HFD-feeding

The HFD-fed (HFF) group showed increased body weight gain after two weeks of HFD-feeding (p < 0.05). The weight gain were noticeable in weeks 2, 4, 6 and 8 weeks of HFD-feeding; with mean percentage weight gain of 7.9%, 22.12%, 27.18% and 31.33% (p < 0.0001) respectively. The HFF group had an elevated 2-hour postprandial blood glucose and insulin levels when compared to the LFD group (p<0.05), indicating impaired glucose metabolism following 8-week HFD-feeding (Table 2). The haematological indices were comparable between the HFD and LFD groups (p>0.05).

**Table 2.** Baseline metabolic and haematological characteristics

| Parameter | LFD | HFD | P value |
| --- | --- | --- | --- |
| Weight (g) | 25.0 ± 2.5 | 26.0 ± 1.9 | 0.43 |
| **Metabolic profile** |  |  |  |
| 2hr postprandial glucose |  |  |  |
| Glucose Levels (mg/dL) | 6.1 (5.4 - 6.9) | 8.7 (8.5 - 9.2) | **0.008** |
| AUC mmol/L*120 min | 636.0 (559.9 - 702.0) | 765.0 (715.5 - 784.5) | **0.032** |
| Insulin conc (μU/ml) | 4.5 (4.4 - 4.6) | 4.8 (4.6 - 8.1) | **0.026** |
| **Haematological parameters** |  |  |  |
| RBC ($10^3\mu$l) | 7.2 ± 0.2 | 6.3 ± 1.0 | 0.14 |
| WBC ($10^3\mu$l) | 4.9 ± 1.0 | 5.5 ± 3.0 | 0.62 |
| MO (%) | 2.0 (2.0 - 2.2) | 2.6 (1.9 - 3.3) | 0.32 |
| LY (%) | 89.2 ± 1.4 | 87.4 ± 3.1 | 0.31 |
| LY # ($10^3\mu$l) | 4.4 ± 0.9 | 4.8 ± 2.6 | 0.72 |
| PLT ($10^3\mu$l) | 701.0 ± 156.3 | 548.8 ± 276.6 | 0.43 |

**LFD:** low fat diet, **HFD:** high fat diet, **AUC:** area under curve, **RBC:** red blood cell, **WBC:** white blood cells, **MO (%):** monocyte percentage, **PLT:** platelet count, **LY (%):** lymphocyte percentage, **LY #:** absolute lymphocyte count.

### Global DNA methylation levels comparison among different cell subsets in prediabetic mice (HFD)

The HFF group had increased %5mC expression on circulating peripheral blood lymphocytes when compared to the LFF group (p=0.049). Notably, T lymphocytes isolated from the HFD group also showed elevated levels of %5mC (p=0.038) when compared to the LFD group. However, in the isolated B lymphocytes the levels of %5mC were comparable between the two diet groups (p = 0.43) (Table 3).

**Table 3.** %5-methyl cytosine levels in HFD-fed compared to LFD-fed mice

| Parameter | LFD | HFD | P value |
| --- | --- | --- | --- |
| CD45<sup>+</sup> Lymphocytes | 5.5 (2.8 – 12.0) | 13.25 (10.3 – 15.9) | **0.049** |
| CD3<sup>+</sup> T cells | 30.9 (14.6 – 43.2) | 68.5 (28.9 – 74.5) | **0.038** |
| CD19<sup>+</sup> B cells | 99.8 ± 0.1 | 99.93 ± 0.03 | 0.432 |

**LFD:** low fat diet, **HFD:** high fat diet. Significant values ( $p < 0.05$ ) shown in boldface.

### Changes in haematological indices in HFD-fed mice treated mice

There were significant differences in the levels of circulating lymphocyte (F _(3,16)_ = 5.44, p = 0.010), monocytes (F_(3,16)_ =3.69, p = 0.033) and platelets (x^2^=9.08, p = 0.028). The post-hoc test showed that the lymphocytes were significantly elevated in the HFD group as compared to the LDA+MET group (p<0.05). In addition, the levels of circulating platelets in the LDA group as compared to the HFD group (Table 4). While no significant differences in the levels of circulating monocytes were observed between the treatment groups (p>0.05) (Table 4).

**Table 4.** %5-methyl cytosine levels in untreated HFD-fed compared to treated HDF-fed mice

| Parameter | HFD | LDA | LDA + MET | CLO | P Value |
| --- | --- | --- | --- | --- | --- |
| **Lymphocytes** | 12.5 ± 12.4 | 6.7 ± 6.4 | 14.1 ± 9.6 | 7.000 ± 4.7 | 0.453 |
| **CD19<sup>+</sup> B cells** | 100.0 (99.9 - 100.0) | 99.7 (99.5 - 100.0) | 99.7 (99.1 - 99.9) | 100.0 (99.7 - 100.0) | 0.087 |
| **CD3<sup>+</sup> T cells** | 41.2 ± 21.9 | 18.8 ± 19.3<sup>a</sup> | 49.88 ± 26.2 | 35.4 ± 28.3<sup>b</sup> | **0.001** |

<sup>a</sup> compared to the HFD group,
<sup>b</sup> compared to the LDA+MET group.
Statistical significance (p < 0.05) is shown in boldface.

### Global DNA methylation levels in HFD-fed compared to treated mice

In order to assess whether DNA methylation profiles are modulated following short-term treatment of 6 weeks, we compared the % 5mC levels across three treatment groups. There were significant changes in the levels of T cells expressing 5mC (F_(4, 20)_ = 6.34, p=0.0016) but not in the total lymphocyte (F_(3, 16)_ = 1.59, p=0,2311) and B cell population (F_(3, 16)_ = 1,472, p=0.2597). The post-hoc test comparing the levels of %5mC expression on T cells showed that LDA significantly lowered levels of %5mC on T cells (p=0.0045). In addition, clopidogrel treatment also lowered the levels of %5mC expressed on T cells when compared to the LDA+MET group (p=0.0051) (Table 4).

## Discussion

This study aimed at measuring the global DNA methylation levels of the major lymphoid cell subsets following short-term high-fat diet feeding. We further assessed whether the levels of DNA methylation are modulated by clopidogrel, low dose aspirin, or low-dose aspirin combined with metformin. The increase in T lymphocyte-specific global DNA methylation in the high-fat diet (HFD) group persisted even during short-term clopidogrel treatment. In fact, our findings showed that elevated DNA methylation levels in T cells following high-fat diet feeding. This may suggest that T cell-specific DNA methylation changes could affect diverse biological signalling that modulate the inflammatory response. Previous studies have reported on an association between obesity and global DNA hypermethylation in lymphocyte subpopulations, including T cells, T cytotoxic cells, and B cells (Jacobsen et al., 2016). Taken together this may suggest that patients with metabolic diseases like T2DM may present with a perturbed epigenetic profile in the subpopulations of circulating lymphocytes. A strong link between inflammation, insulin resistance, and epigenetic modifications has been reported (Zhao et al., 2012; Van Otterdijk et al., 2017). T cells play a crucial role in the initiation as well as maintenance of inflammation in adipose tissue, through persistent macrophage recruitment which may as a consequence promote the development of insulin resistance (Winer et al., 2011; Winer et al., 2012; Nishimura et al., 2009; DeFuria, A. C. Belkina, et al., 2013).

Epigenetic modification, precisely DNA methylation, may affect clopidogrel response (Zhang et al., 2017). Age and body mass index (BMI), which are some of the risk factors for type 2 diabetes and metabolic syndrome, have also been reported to be significantly associated with clopidogrel response (Cuisset et al., 2009; Khalil et al., 2016). It has also been reported that diabetic patients exhibit a poor antiplatelet effect upon treatment with clopidogrel when compared to non-diabetics (Angiolillo et al., 2005; Serebruany et al., 2008), a phenomenon that is yet to be elucidated.

Although low-dose aspirin and clopidogrel have been used as dual therapy in the primary prevention of cardiovascular events in patients with T2DM. A substantial incidence of major adverse cardiac events (MACE) has been reported. Exploration of the role and association of T cell-specific DNA methylation with clopidogrel action could help in the elucidation of the possible factors associated with the varied patient responses (Khalil et al., 2016). In fact, Garcia-Calzon *et al.*, reported that DNA methylation on metformin transporter genes in the human liver, which differed according to anti-diabetic drug that was administered (Garcia-Calzon et al., 2017). In our study, LDA significantly decreased the levels of T cell specific global DNA methylation in a prediabetic state. Surprisingly, no synergetic modulation of T cell global DNA methylation was observed in our study. These novel findings could suggest potential insight in the variable responses to anti-inflammatory drugs amongst patients living with T2DM. Persistent T cell activation that is initiated at the prediabetic phase may persist during treatment and lead to increased thrombotic risk.

The current study was limited to the major cell lymphocyte lineages and no T cell subtyping was performed to delineate whether differences in the T cell subsets exist. Activated T cells have been implicated in coronary artery disease remain one of the macrovascular complications associated with type 2 diabetes mellitus. In addition, an epigenome-wide study revealed that hypomethylation within the tumour necrosis factor receptor-associated factor 3 (TRAF3) gene was associated with increased platelet aggregation and vascular recurrence in ischemic stroke patients who were under clopidogrel treatment (Gallego-Fabrega et al., 2016). Moreover higher TRAF3 expression due to decreased methylation may lead to an increase in the CD40 signal pathway (Song et al., 2011; Kuijpers et al., 2015). CD40 is involved in the co-stimulation and activation of T cells (Song et al., 2011). It remains unclear whether hypermythylated T cells retain the functional capacity and whether in this may affect immunological responses in patients living with T2DM.

## Conclusion

T cells are involved in the initial perturbation of DNA methylation profiles in the pathogenesis of inflammation, insulin resistance and subsequently type 2 diabetes. Low-dose aspirin is effective in modulating T cell-specific global methylation, whereas clopidogrel showed no modulatory effect on the DNA methylation profile following a short term high fat diet feeding. This may suggest that the early changes in T cell DNA methylation profiles are mediated by inflammation and may be reversed by using low-dose aspirin.

## Authors’ contributions

TM and BBN conceived the idea and design of the study as well as results analysis. ZM and PVD helped draft the article. All authors wrote and approved the final manuscript.

## Ethics approval

The study was approved by the University of KwaZulu Natal’s Animal Research Ethics Committee (AREC) with the ethics number: AREC/086/016.

## Acknowledgements

The University of KwaZulu-Natal (UKZN) Biomedical Resource Unit (BRU) is acknowledged for the assistance with mice laboratory procedures and animal housing facilities. Furthermore, the UKZN Department of Human Physiology, College of Health Sciences (CHS) is acknowledged for providing access to the flow cytometry analysis facility.

